# Identification of genes associated with abiotic stress tolerance in sweetpotato using weighted gene co-expression network analysis

**DOI:** 10.1101/2023.01.30.526063

**Authors:** Mercy Kitavi, Dorcus C. Gemenet, Joshua C. Wood, John P. Hamilton, Shan Wu, Zhangjun Fei, Awais Khan, C. Robin Buell

## Abstract

Sweetpotato, *Ipomoea batatas* (L.), a key food security crop, is negatively impacted by heat, drought, and salinity stress. We exposed the orange-fleshed cultivar ‘Beauregard’ to 24 and 48 hours of heat and salt stresses to identify differentially expressed genes (DEGs) in leaves. Analysis revealed both shared and unique sets of up-regulated (650 for heat; 287 for salt) and down-regulated (1,249 for heat; 793 for salt) DEGs suggesting common, yet stress-specific transcriptional responses to these two abiotic stressors. Gene Ontology analysis of downregulated DEGs common to both heat and salt stress revealed enrichment of terms associated with ‘cell population proliferation’ suggestive of an impact on the cell cycle by the heat stress. To identify shared and unique gene coexpression networks under multiple abiotic stress conditions, weighted gene co-expression network analysis was performed using gene expression profiles from heat, salt, and drought stress treated ‘Beauregard’ leaves yielding 18 coexpression modules. One module was enriched for ‘response to water deprivation’, ‘response to abscisic acid’, and ‘nitrate transport’ indicating synergetic crosstalk between nitrogen, water and phytohormones with genes encoding osmotin, cell expansion, and cell wall modification proteins present as key hub genes in this drought-associated module. This research lays the background for future research in mediating abiotic stress tolerance in sweetpotato.

## 1. Introduction

Crops are exposed to abiotic and biotic stress which can lead to adverse effects on growth and productivity (Hussain et al., 2019). Changing climate conditions including extreme temperature and water deficit conditions have the potential to substantially limit crop growth and productivity (Dahal et al., 2019; Hussain et al., 2019; Zhou et al., 2017). These production limitations, coupled with the need to expand feed and food production to meet a growing world population, will most likely require expansion of production areas to marginal agricultural lands. Understanding how plants perceive abiotic stress signals and adapt to adverse environmental conditions can facilitate development of improved cultivars essential for global food security.

Sweetpotato, *Ipomoea batatas* (L.) Lam., which originated in Central America, is a widely cultivated crop with a world production in 2019 of 91.8 metric tons (FAOSTAT, 2021). Sweetpotato root flesh can be white, cream, yellow, orange, or purple in color and are excellent source of vitamins (C, E, K and several B vitamins) and a major caloric source for sub-Saharan Africa (Bovell-Benjamin et al., 2009; Low et al., 2017). However, only the orange fleshed sweetpotato varieties are rich in beta-carotene, which is converted into vitamin A, an essential vitamin for a strong immune system, healthy skin, as well as vision and eye health (Low and Thiele, 2020). Currently, efforts to address vitamin A in sub-Saharan Africa are focused on use of orange-fleshed sweetpotato (Low et al., 2017) and Beauregard, an orange-fleshed cultivar with low dry matter (Lau et al., 2018; Rolston et al., 1987), has played a critical role in the improvement of African sweetpotato cultivars with increased β-carotene *via* introgression of *Orange* (*Or)* alleles through breeding (Gemenet et al., 2020). Beauregard has been used in both direct and recurrent crosses for studying of agronomic traits such as root -knot nematode resistance, drought tolerance and understanding of the relationship between starch and beta-carotene in elite breeding lines (Cervantes-Flores et al., 2011; Gemenet et al., 2020; Lau et al., 2018; Mollinari et al., 2020).

Plants have evolved interconnected regulatory pathways enabling response(s) to abiotic and abiotic stress. Understanding these mechanisms can aid in understanding and improving stress tolerance and the potential to recover yield under stress (Atkinson and Urwin, 2012; Bashir et al., 2019; Gull et al., 2019). Furthermore, transcriptional regulatory networks in *Arabidopsis thaliana* have revealed some of the underlying molecular, cellular, physiological, and biochemical bases of abiotic stress responses (Barah et al., 2016; Berens et al., 2019; Nakashima et al., 2009; Yamada et al., 2010). This includes physiological changes such as altering stomatal pore apertures thereby enabling optimized CO_2_ uptake while minimizing water loss (Nilson and Assmann, 2007) as well as minimizing oxidative damage from the production of reactive oxygen species (ROS) (Arbona et al., 2017).

Phytohormones are crucial integrators of the adaptive mechanisms of stress responses. Abscisic acid (ABA)-dependent and ABA-independent pathways have been described in the response to drought stress (Yamaguchi-Shinozaki and Shinozaki, 2006). Plant leaves respond to water deficit by increasing ABA biosynthesis, transport and accumulation thereby triggering stomatal closure that decreases gas exchange rate, respiration, and photosynthetic activity (Osakabe et al., 2014; Yamaguchi-Shinozaki and Shinozaki, 2006). ABA action not only targets guard cells and the induction of stomatal closure but also systemically signals to adapt to water limitations. Due to its induction by various stresses, ABA is considered a plant stress hormone (Swamy and Smith, 1999; Tuteja, 2007). Several transcription factors (TFs) are known to regulate the expression of abiotic stress-responsive genes either via the ABA-dependent or ABA-independent pathway (Xiong et al., 2002).

Gene expression profiling experiments can provide insight into plant responses to environmental stressors (Lau et al., 2018; González-Schain et al., 2019, 2016; Mangrauthia et al., 2017; Frey et al., 2015; Baidyussen et al., 2020; Rensink et al., 2005; Gong et al., 2015; Zhang et al., 2017; Arisha et al., 2020b; Tang et al., 2020; Kumar et al., 2021; Li et al., 2019). Access to large-scale gene expression profiling datasets from a wide range of species under diverse abiotic stress conditions has revealed not only stress-specific responses but also shared or common responses to multiple abiotic stress conditions. Weighted gene co-expression network analysis (WGCNA) clusters genes into co-expression networks based on the connectivity of the gene expression (Zhang and Horvath, 2005; Langfelder and Horvath, 2008) in which genes with similar expression patterns are grouped into the same module (clusters of highly interconnected genes) suuggestive of similar functions and/or potentially common biological regulatory roles (Zhou et al., 2018). This method has been successfully used to identify modules associated with biological processes in different plant species (Du et al., 2017; Greenham et al., 2017; Shaik and Ramakrishna, 2013; Hoopes et al., 2019; Wang et al., 2020). Within these modules are hub nodes (genes) which are highly connected, central to the network’s architecture, and are hypothesized to have an influential role in regulating network structure (Barabási and Oltvai, 2004; Albert et al., 2000). Gene connectivity has been used for identifying hubs as well as identifying differentially connected genes (Arnatkeviciute et al., 2021; Liu et al., 2019b; Panahi and Hejazi, 2021).

Generation of the reference genome sequence of the wild diploid sweetpotato relative, *Ipomoea trifida* (Kunth) G.Don (Wu et al., 2018), has permitted transcriptome analysis of cultivated hexaploid sweetpotato (Gemenet et al., 2020; Bednarek et al., 2021; Suematsu et al., 2020). With respect to abiotic stress, transcriptomic responses of sweetpotato under simulated drought conditions (Arisha et al., 2020b; Lau et al., 2018), heat stress (Arisha et al., 2020a), and salt stress (Luo et al., 2017; Meng et al., 2020; Yang et al., 2020b) have been reported. Here, we examined gene-expression patterns following heat and salt stress treatments in leaves of the orange-fleshed sweetpotato cultivar Beauregard. We identified shared and stress-specific differential gene expression over a 48 hr time course. A broader view of abiotic stress in cultivated sweetpotato was established by constructing gene co-expression networks using gene expression datasets from this study combined with a previously published simulated drought stress dataset (Lau et al., 2018).

## 2. Materials and methods

### 2.1 Experimental design and stress treatments

*In vitro* plantlets of cultivar ‘Beauregard’ were grown on sterilized MPB solid medium (Murashige and Skoog salts, 3% sucrose, 2 mg/L calcium pantothenate, 100 mg/L L-arginine, 200 mg/L ascorbic acid, 20 mg/L putrescine-HCl, 10 mg/L GA_3_, 0.3% Phytagel, pH 5.7) at 28°C/20°C day/night temperature with 12-hr light and 12-hr dark conditions, under 3000 lx and relative humidity of 70%. After 14 days, plants were transferred to MPB liquid media and left to acclimatize to the media for 7 days, after which either stress or control treatments were imposed. For salinity stress, the medium was amended with 150 mM NaCl with the temperature kept at 28°C/20°C day/night; for heat stress, the plants were grown at 40°C/32°C day/night. For control treatments, the liquid media was also replaced but no stress treatment was applied. Leaf samples were collected at 24 hr and 48 hr after stress (HAS) from salt and heat treated plants for stress, and no stress treated plants for control.

### 2.2 Library construction and RNA sequencing

Leaf samples were collected in three biological replicates per treatment and timepoint. The TRIzol (Invitrogen) method was used to extract RNA and sequencing libraries were constructed using the protocol described in Zhong et al. (2011) and sequencing done on an Illumina HiSeq 2500 platform. Raw RNA-Seq reads were filtered and trimmed using Cutadapt v2.10 (Martin, 2011) to remove adapter sequence, low-quality bases (quality score < 30), retaining only reads with a minimum read length of 70 nt. Libraries sequenced to 151 nt were clipped to 101nt using Cutadapt v2.10 (Martin, 2011) to equalize read length. Filtered clean reads were assessed for quality using FastQC v0.11.8 (https://www.bioinformatics.babra-ham.ac.uk/projects/fastqc) and viewed with MultiQC v1.9 (Ewels et al., 2016). Cleaned reads were aligned to the diploid reference genome *I. trifida* (NSP306; v3; http://sweetpotato.uga.edu/) (Wu et al., 2018) using HISAT2 v2.2.1 (Kim et al., 2019) with the options: --phred33 --min-intronlen 20 --max-intronlen 5000 --rna-strandness R --dta-cufflinks. Aligned files were sorted and merged using the samtools sort and merge options, respectively, in SAMTools v1.10 (Li et al., 2009). RNA-seq data from Beauregard plants exposed to polyethylene glycol (PEG) to mimic drought were obtained from Lau et al. (2018).

Expression abundances (fragments per kilobase of exon model per million mapped fragments; FPKM) for each gene model were determined based on the RNA-Seq read alignments using Cufflinks v2.2.2 (Trapnell et al., 2010) with the parameters --multi -read -correct, --min - intron -length 10, --max -intron -length 5000, --library -type fr -first strand. Read counts were determined using HTSeq v0.12.3 (Anders et al., 2014) with the parameters --stranded = reverse -- minaqual = 10 -type = exon --mode = union. For sample-level quality controls, the relationship between the biological replicates was visualized using Pearson Correlation Coefficients (PCC) of gene expression abundances. Some libraries were sequenced more than once (technical replicates) and PCCs were checked prior to merging technical replicates; all technical replicates had a PCC > 0.97. To assess batch effects of the experiments, the regularized-logarithm transformation (rlog) of expression count data was used to generate a Principal Component Analysis (PCA) and heat map in R v4.1 (https://cran.r-project.org/).

### 2.3 Differential gene expression

Using the unique combination of treatment and time -point as a single factor, gene counts were used to detect differentially expressed genes (DEGs) with DESeq2 v1.28.1 (Love et al., 2014) between stress treatments and their respective controls in R v4.1. To detect genes that responded differentially to each treatment, contrasts were made between treatment and control at each time point using the model, design = ∼ treatment + time performed in DESeq2. Genes were filtered at an adjusted *p* -value (padj) cut -off < 0.05 and log_2_FoldChange (LFC) of 2.0. For visualization, “lfcShrink” function was used to account for low expression levels and MA plots were generated for each treatment and timepoint. Genes with a fold-change between 2.0 for upregulated and -2.0 for downregulated genes with an adjusted *p*-value cutoff of 0.05 were deemed DEGs. Gene Ontology (GO) enrichment tests were performed using topGO v2.50.0 in R v4.1 (Alexa and Rahnenfuhrer, 2021). *P*-values < 0.05 from Fisher’s Exact Tests using the weighted model was used as the test for significance (Benjamini and Hochberg, 1995).

### 2.4 Clustering and construction of gene co-expression networks

Differentially expressed genes were clustered using the unsupervised K-means clustering method with parameters: max_itr = 10000 and n_clust = 25 in R (Oyelade et al., 2016). Co-expression of genes was further explored using WGCNA in R v4.1.1 (Langfelder and Horvath, 2008). The expression abundances (FPKMs) of all gene isoforms (44,158) were used and filtered to 14,138 genes by removing all features whose counts were consistently low with a count of less than 5 in all the samples, (x < 5) in heat, salt, and drought treated samples.

Samples were initially clustered using the FlashClust tool to analyze sample height and detect and remove outliers. A soft thresholding power of 20 was chosen for subsequent co-expression module construction following application of the scale-free topology criterion described by Barabási (2009). Based on the topological overlap-based dissimilarity measure (Zhang and Horvath, 2005), genes were hierarchically clustered, and a dendrogram was used for module detection using the dynamic tree cut method (mergeCutHeight = 0.9, minModuleSize = 35). Modules were identified as gene sets with high topology overlap measure (TOM) generated by the adjacency and correlation matrices of gene expression profiles. Genes that did not fit in any modules were discarded from further analyses (gray module). Gene ontology enrichment of module genes was performed using topGO package; a *p-*value or FDR < 0.05 was used to determine the significant enrichment (Alexa and Rahnenfuhrer, 2021).

Gene connectivity based on the edge weight (ranging from 0 to 1) was determined by TOM; the weights across all edges of a node were summed and used to define the level of connectivity, and nodes with high connectivity were considered hub genes. Intramodular connectivity measure, *kWithin*, or connectivity of a particular gene to all other genes within its same module was used to define hub genes in each module. *kTotal* was used to measure the connectivity of a gene to all other genes regardless of module (*kOut* is kTotal-kWithin, and *kDiff* is kWithin-kOut) (Rhead et al., 2020).

### 2.5 Gene orthology

OrthoFinder v2.5.4 (Emms and Kelly, 2015) was run with the longest peptide isoform of each *I. trifida* predicted peptide (http://sweetpotato.uga.edu/) with the *A. thaliana* TAIR10 (https://www.arabidopsis.org) and *Oryza sativa* v7 predicted proteome (Ouyang et al., 2007) obtained from Phytozome v12.1.5 (Goodstein et al., 2012). Transcription factors in *I. trifida* were obtained from iTAK (Zheng et al., 2016).

### 2.6 Data availability

Raw sequence reads have been deposited in the National Center for Biotechnology Information Sequence Read Archive BioProject PRJNA834099 and PRJNA834095.

## 3 Results and Discussion

### 3.1 Dynamic expression patterns of differentially expressed genes following heat and salt stress in cultivated sweetpotato

A total of 55 Beauregard RNA-seq samples (technical and biological replicates) from the control, salt stress, and heat stress were processed revealing an overall alignment rate to the *I. trifida* reference genome between 72.12% and 85.2% (**Table S1**), consistent with previous analyses in which transcript reads from sweetpotato were aligned to *I. trifida*, a close diploid ancestor of the cultivated hexaploid *I. batatas* (Lau et al., 2018; Wu et al., 2018). Technical replicates were merged post-alignment resulting in a total of 24 control and stress samples. Correlations between biological replicates were examined by generating pairwise PCCs for gene expression abundances (FPKMs); an average PCC value of 0.9985 and a minimum PCC of 0.9912 was observed (**Table S2**) suggesting high reproducibility and quality of the datasets.

A PCA (**Figure 1a)** was plotted to assess the relationships of the biological replicates and the extent of gene expression variation explained by the experimental variable. The tight clustering of biological replicates revealed that the largest variation of gene expression observed was due to the experimental conditions (heat vs salt and stress vs control). PC1 contributed 64% whereas PC2 contributed 17%. Limited separation of samples based on treatment timepoint was observed suggesting that gene expression within the treatment or control is more similar than between treatments or controls.

**Figure 1.**
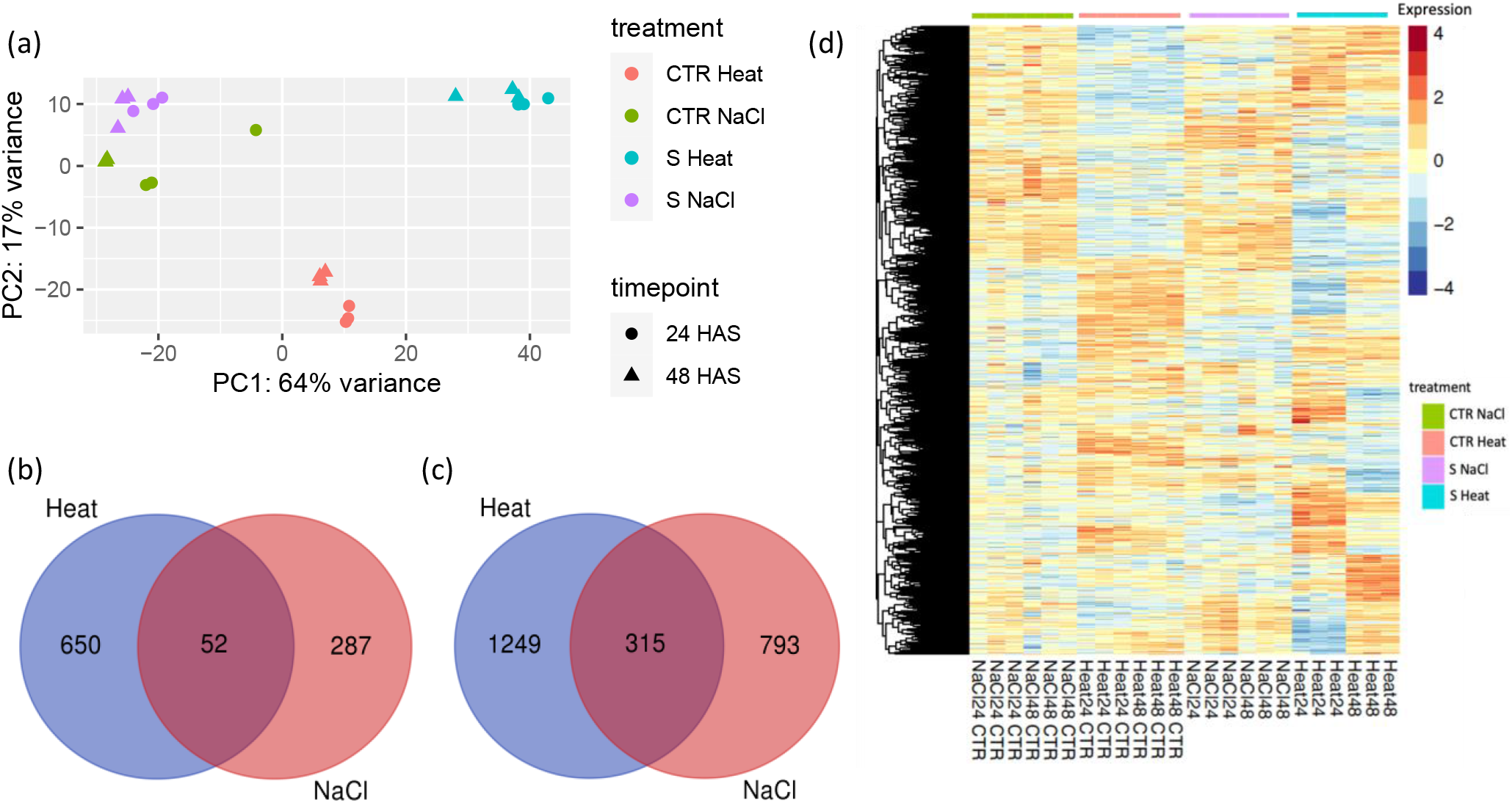
Assessment of inter- and intragroup variability. (a) Principal component analysis plot displaying all replicates along principal components 1 and 2 explaining 64% and 17%, respectively, of the variance in gene expression. Replicate samples from the same group cluster together, while samples from different groups form separate clusters indicating that the differences between groups are larger than those within groups. Venn diagrams indicating the number of differentially expressed genes (DEGs); (b) upregulated DEGs and (c) downregulated DEGs. The intersection shows genes commonly regulated by both stress conditions. (d) Heatmap visualizing clustering patterns of 3,346 DEGs (*p*-value < 0.05) in salt (150 mM NaCl) and heat (40°C) treated Beauregard leaf tissue suggesting stress specific and shared gene expression under these two stress conditions. S and CTR denotes stress and control samples, respectively

Analysis of DEGs revealed unique and shared sets of up- and down-regulated genes following heat and salt stress at the two timepoints (24 and 48 HAS; **Table S3**). A total of 433 genes were upregulated by heat at 24 HAS while 502 genes were up-regulated at 48 HAS; 236 genes overlapped between the two timepoints leaving 702 unique genes. Under salt stress, 288 genes were upregulated at 24HAS, and 182 genes were up-regulated at 48HAS with 72 genes shared between the two timepoints. Treatment of sweetpotato leaves with heat caused downregulation of 974 genes at 24 HAS and 1146 genes at 48 HAS (556 shared genes between timepoints). Salt resulted in down-regulation of 955 genes at 24 HAS and 426 at 48 HAS, with 272 genes were down regulated in both time points. With the substantial overlap in genes differentially regulated in these two timepoints, we merged the DEGs at both timepoints for each stress and performed downstream analyses with the non-redundant set of DEGs for each stress.

Comparing the salt and heat treatments, the differentially upregulated genes totaled 989; 650 unique to heat stress, 287 unique to salt stress and 52 genes common to the two stress conditions (**Figure 1b, Tables S4, S5, S6)**. With respect to down-regulation, 2,357 genes were differentially expressed in these two stress conditions with1,249 were unique to heat, 793 unique to salt stress, and 315 genes downregulated in both stress treatments (**Figure 1c; Tables S7, S8, S9)**. The partitioning of DEGs unique to each stress is consistent with previous reports on the specificity of plant acclimations to individual stressors (Koch and Guillaume, 2020; Krasensky and Jonak, 2012) and visible in hierarchical clustering of DEGs (**Figure 1d**) in which shared as well as distinct clustering patterns are apparent.

Abiotic stressors including drought, cold, salt, heat, oxidative stress, and heavy metal toxicity strongly influence photosynthesis (Huang et al., 2019; Yang et al., 2020a; Cornic, 2000). A total of 315 genes were differentially downregulated genes overlapping between heat and salt stress. The GO term ‘chlorophyll biosynthetic process’ (GO:0015995; *p-*value 0.00119) (**Figure 2**) was enriched among the 315 genes and contained 12 photosynthetic genes including cycloartenol synthase (*CAS1*) which affects chlorophyll and carotenoid biosynthetic pigments in plants (Babiychuk et al., 2008; Luo et al., 2019). Various experiments have demonstrated a sharp decline in chlorophyll content (chlorosis) and decrease in expression of some essential enzymes under stress (Allakhverdiev et al., 2008; Rossi et al., 2017). Destruction of the chloroplast ultrastructure by stress leads to a decrease in chlorophyll and ultimately lower photosynthetic activity (Hamani et al., 2020; Sidhu et al., 2017) explaining the downregulation of photosynthesis-related genes (**Figure 2**). Moreover, the decrease in the chlorophyll content can also be caused by a decrease in the stomata aperture to limit water losses by evaporation and increased resistance to the entry of atmospheric CO2 necessary for photosynthesis..

**Figure 2.**
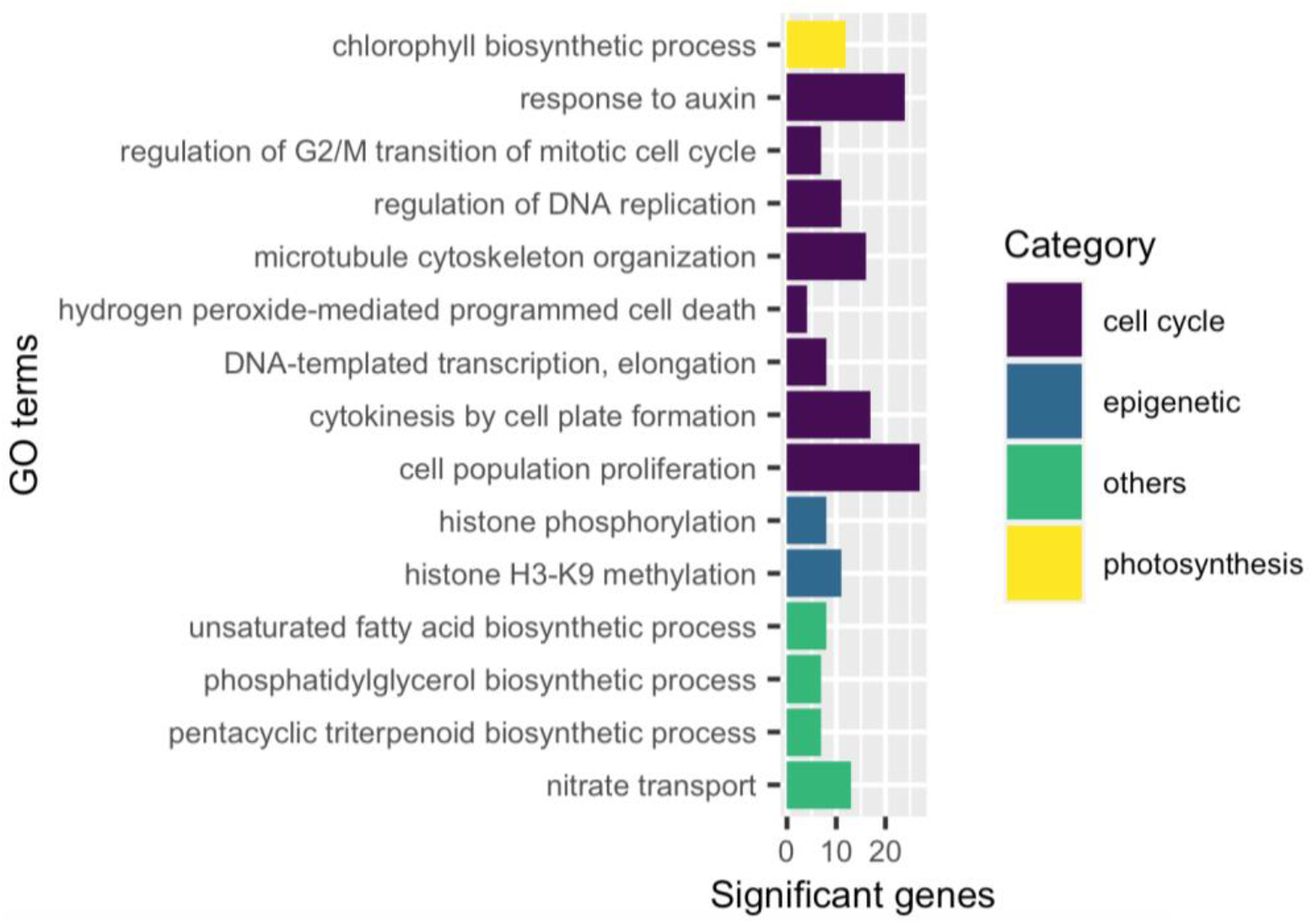
A graphical representation of gene ontology enrichment and gene annotation terms of the genes differentially down-regulated (DEGs; LFC < - 2.0; *p-*value < 0.05) by both heat and salt stress in Beauregard leaf samples.

Organ growth is divided into cell proliferation (increase in cell number) and the cell expansion stage (expansion to the final size). Arrest of the cell cycle in response to stress enables organisms to survive under fluctuating environmental conditions or DNA damage thereby allowing DNA repair to occur before DNA replication or mitosis resumes (Hu et al., 2016). DEGs that were downregulated by both heat and salt (**Figure 2**; **Table S9)** were enriched in GO terms associated with ‘cell population proliferation’ (GO:0008283392, *p-*value 4.30E-13) and included genes encoding kinases and other genes previously reported to participate in plant cell cycle control in *A. thaliana* (Stals et al., 2000; Bertoni, 2018). During plant cytokinesis, Golgi/trans-Golgi network-derived vesicles are targeted to the plane of cell division where they fuse to form the cell plate. An ortholog of Qa-SNARE (itf13g15140), a specialized syntaxin named KNOLLE (SYNTAXIN OF PLANTS 111, SYP111), is required for cytokinesis membrane fusion (Boutté et al., 2010; Lauber et al., 1997; Lukowitz et al., 1996; Touihri et al., 2011; Waizenegger et al., 2000) and exhibited significant down-regulation following 48 hr of heat (LFC = -6.8099, *p-*value 4.54E-14) and 24 hr salt (LFC = -2.1419, *p-*value 5.71E-05) stress (**Table S9**). Perturbation of cell cycle progression impacts the chromatin state and GO enrichment of terms associated with ‘histone phosphorylation’ (GO:0016572; *p*-value 0.001) and ‘histone H3-K9 methylation’ (GO:0051567; *p-*value 0.013) were observed among the heat and salt common downregulated DEGs (**Figure 2**). Chromatin-associated genes down-regulated by heat and salt include two HISTONE 3.3 genes (itf09g16990: heat 48 hr, LFC = -2.8211, *p-*value 2.50E-06; salt 24 hr, LFC = -2.0186, *p*-value 0.0005 and itf06g11890: heat, 48 hr, LFC = -2.8828, *p-*value 7.21E-05; salt, 24 hr, LFC = -2.4259, *p-*value 0.0005) (**Table S9**). In addition, multiple Small Auxin Upregulated RNA (SAUR)-like auxin responsive protein family genes are among the 315 common down-regulated DEGs under heat and salt stress (**Table S9**). SAUR-like auxin responsive protein family genes have been shown to regulate adaptive growth and are generally down-regulated in response to abiotic stress.

### 3.2 Heat stress in sweetpotato triggers a canonical heat shock response

Differentially expressed genes following heat stress were significantly enriched in 115 GO terms sorted into three categories, *Biological process* (BP), *Cellular component* (CC), and *Molecular function* (MF) (*p*-value **≤** 0.05; **Table S10**), across the two time-points. Similar to that observed in Arabidopsis heat stress (Grinevich et al., 2019), heat-related *Biological Process* GO terms such as ‘response to hydrogen peroxide’ (GO:0042542: *p*-value; 7.60E-28), ‘response to high light intensity’ (GO:0009644; *p*-value: 4.90E-27), ‘protein folding’ (GO:0006457; *p*-value: 1.70E-25), and ‘response to heat’ (GO:0009408: *p*-value; 4.60E-15) were significantly enriched (**Figure 3a)** among the DEGs up-regulated following heat stress suggesting that even brief exposure to 40 °C resulted in perceived stress in *I. batatas* leaves.

**Figure 3.**
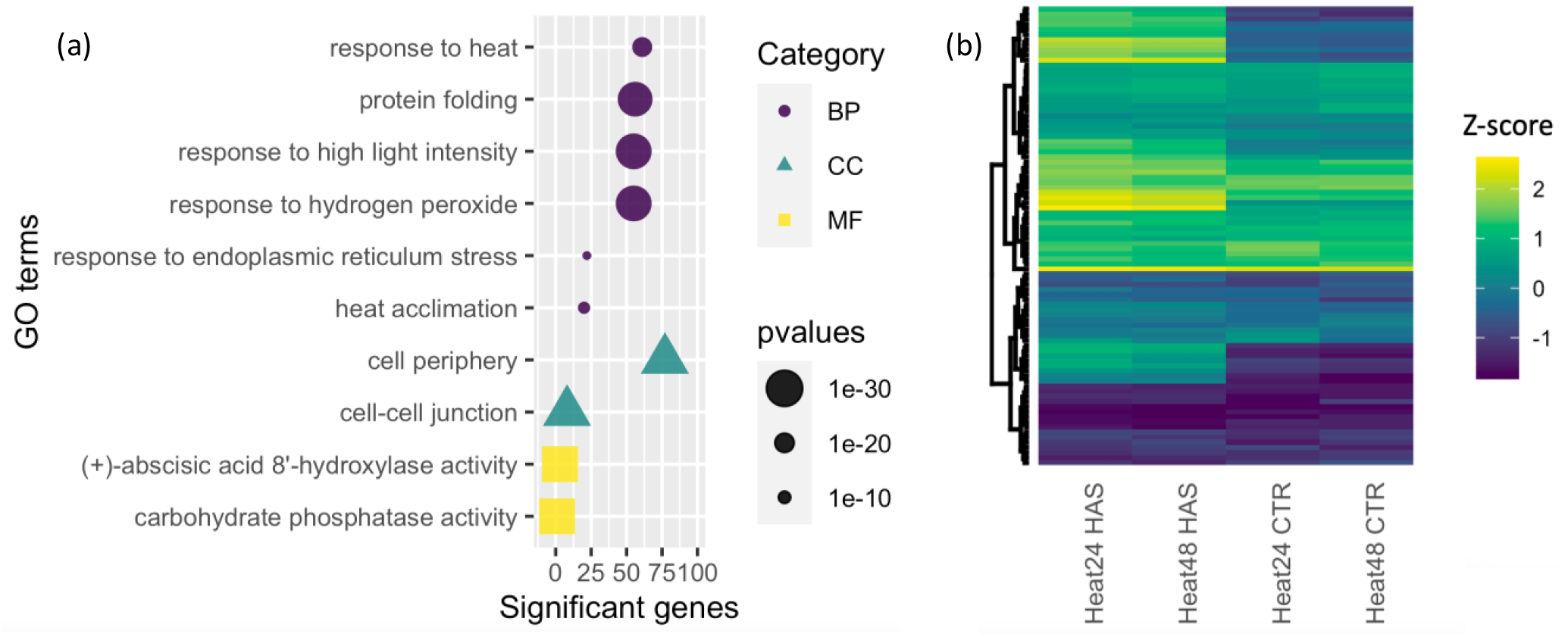
Effects of heat stress on Beauregard leaf. (a) Gene Ontology (GO) terms enriched for the 702 upregulated DEGs following heat stress. GO terms are sorted into three categories; *Biological process;* BP, *Cellular component;* CC, and *Molecular function;* MF. (b) Heatmap of the expression (Z-score) of heat shock proteins (HSPs) and heat shock transcription factors (HSFs) indicating their regulation (down-or up-regulated based on the scale color) in Beauregard leaves following heat stress.

Activation of transcriptional cascades following heat stress is common in many plant species. Following heat stress in Arabidopsis, 21 heat shock transcription factors (HSFs) and >2500 genes were differentially expressed (Busch et al., 2005). We identified 94 *I. trifida* orthologs that encode heat shock-related proteins (HSPs) and HSFs (**Table S11, Figure 3b**).

Following heat stress, 28 HSP and 4 HSFs were differentially expressed with upregulation primarily occurring at 48 HAS rather than 24 HAS. A delay in the induction of heat shock proteins suggests that sweetpotato has a prolonged temperature threshold for the canonical heat shock response induction. This parallels the response in tepary bean (Frijol Bayol) relative to common bean, in which tepary bean which is adapted to hotter climates had a higher temperature threshold for induction of the canonical heat shock response (Moghaddam et al., 2021).

Arabidopsis chloroplast HSP mutants are sensitive to heat shock and exhibit variegated cotyledons, malformed leaves, growth retardation, and impaired root growth following heat stress (Schroda et al., 1999; Lee and Schöffl, 1996). In 48 hr heat stressed Beauregard leaves, the chloroplast-localized small heat shock protein HSP21 ortholog (itf00g29810, 48 hr, Log_2_FC = 11.4101, *p*-value 7.87E-20) was the highest and most significantly upregulated gene (**Table S4)**, consistent with observations in Arabidopsis where HSP21 is involved in heat acclimation and was hyper-induced in response to heat stress in Arabidopsis (Kim et al., 2014). CHLOROPLAST HEAT SHOCK PROTEIN 70-1 (itf04g21320, 48 hr, Log_2_FC = 5.1012; *p*-value 2.07E-117) was also significantly upregulated following heat stress (**Table S4)**, yet the three paralogs of itf04g21320 [itf10g21850 (cpHSP 70-1), itf01g02720 (cpHSP 70-1), itf09g04300 (cpHSP 70-2)] were not differentially expressed suggesting neo-functionalization at the expression level in this gene family. Small heat stress proteins were the most abundant of the upregulated heat shock proteins in this study and are the most heat-responsive in plants due to their dramatic induction and prevention of the aggregation of heat-labile proteins that stabilize lipids at the plasma membrane while the HSP chaperones prevent and repair protein misfolding and aggregation to reduce cell damage (Guihur et al., 2021).

Heat shock transcription factors (HSF) play a critical role in the response to heat stress (Ohama et al., 2017) and HSFs have been identified in *Arabidopsis* (Nover et al., 2001), tomato, rice, maize (Scharf et al., 2012), wheat (Xue et al., 2014), apple (Giorno et al., 2012), poplar (Liu et al., 2019a; Zhang et al., 2015), desert poplar (Zhang et al., 2016), pear (Qiao et al., 2015), tea (Liu et al., 2016), and grape (Liu et al., 2018). In this study, HSFA6B gene (itf04g25720, 48 hr, LFC = 3.3552, *p*-value 0.0004, **Table S4**) was up-regulated following heat stress. Simultaneous editing of *HSFA6a* and *HSFA6b* in Arabidopsis caused a reduction in reactive oxygen species (ROS) accumulation and increased expression of abiotic stress and ABA-responsive genes, including those involved in ROS level regulation suggesting their involvement in abiotic stress tolerance through the regulation of ROS homeostasis in plants (Wenjing et al., 2020). HSFA2 genes are necessary for the maintenance of the acquired thermotolerance (Lämke et al., 2016) and play a key role in the initiation of plant heat stress responses in Arabidopsis (Ohama et al., 2017). HSFA2 (itf12g20490, 48 hr, LFC = 2.2994, *p*-value 3.40E-05, **Table S4**) was up-regulated upon heat stress. Increased HSFA2 levels in sweetpotato corroborate findings from Arabidopsis in which overexpression of the Arabidopsis HsfA2 gene not only increased thermotolerance but also salt/osmotic stress tolerance compared to the wild-type (Ogawa et al., 2007).

Both by enzymatic and nonenzymatic pathways causes H_2_O_2_ production in plants under natural and stressful conditions and is associated with plant development and growth. Accumulation of ROS and development of secondary stress responses under elevated temperature is because of heat on the membrane fluidity (Driedonks et al., 2015; Han et al., 2019; Mittler et al., 2012; Xiong et al., 2021). Although H_2_O_2_ is toxic at high concentrations, heat stress-induced H_2_O_2_ is required for the effective expression of HSP genes in Arabidopsis (Volkov et al., 2006; Wang et al., 2014). HSFs are postulated to act as H_2_O_2_ sensors in the plants (Davletova et al., 2005; Miller and Mittler, 2006). Consistent with this hypothesis, 58 genes were enriched for the GO term ‘response to hydrogen peroxide’ (GO:0042542, *p*-value 7.60E-28) including the HEAT SHOCK TRANSCRIPTION FACTOR B2A; HSFB2A (itf13g03340; 24 hr, LFC = 3.3064, *p*-value 8.38E-11) following heat stress in sweetpotato.

We observed enrichment of GO terms in genes down-regulated in response to heat stress associated with the cell cycle including ‘cell population proliferation’ (GO:0008283, *p*-value 3.20E-20), ‘cytokinesis by cell plate formation’ (GO:0000911, *p*-value 3.00E-09), ‘G2/M transition of mitotic’ (GO:0010389, *p*-value 1.90E-07), ‘DNA replication’ (GO:0006275, *p*-value 1.40E-11), and ‘DNA replication initiation’ (GO:0006270, *p*-value 5.50E-10) (**Table S10)**. Increased cellular temperatures cause protein denaturation interrupting critical cellular processes and resulting in apoptosis and cell death (Gao et al., 2015; Gu et al., 2014; Matsuki et al., 2003). The cell cycle consists of G1, S, G2, and M stages with two major checkpoints at the G1/S checkpoint and G2/M. Cyclins in combination with cyclin-dependent kinases (CDKs) drive the cell cycle. Abiotic stress induces signaling molecules that suppress the activities of CDKs via controlling the expression level of cyclins or regulating the posttranslational modification of CDKs therefore arresting or even exiting the cell cycle. Transient arrest of cell division at G1/S and G2/M transitions was reported in Arabidopsis following a short-term (acute) exposure to moderate heat shock, (Kühl and Rensing, 2000; Harashima et al., 2013) and in tobacco BY2 cells under a mild heat stress (Jang et al., 2005). Moghaddam et al. (2021) observed down-regulation of cellular process and cell cycle arrest following exposure to short moderate heat stress in heat tolerant tepary bean (*Phaseolus acutifolius*) but not heat sensitive common bean (*Phaseolus vulgaris*) suggesting adaptive responses to heat stress between these two *Phaseolus* species. In heat stressed sweetpotato, the cyclin-dependent kinase B2;2 (CDKB2;2 itf07g23920, LFC = -2.17237, *p*-value 1.24E-06) gene involved in cytokinesis and regulation of the G2/M transition of the mitotic cell cycle was downregulated.

### 3.3 Gene expression profiles following salt stress

High levels of salt lead to ion toxicity, hyperosmotic stress, and secondary stresses (Chakraborty et al., 2018; Zhu, 2002). Na^+^, if accumulated in the cytoplasm, can be toxic to living cells adversely affecting K^+^ nutrition and vital plant physiological mechanisms including cytosolic enzymes, photosynthesis, and metabolism (Chakraborty et al., 2018; Shabala and Cuin, 2008). Salt stress impacts plant growth and development by reducing plant water potential, altering nutrient uptake, and increasing the accumulation of toxic ions (Shrivastava and Kumar, 2014). Among the enriched GO terms for genes differentially upregulated by salt stress were those associated with ‘response to water deprivation’ (GO:0009414, *p*-value 2.90E-05) and ‘hyperosmotic salinity response’ (**Figure 4, Table S10**). The homeobox gene (ATHB7, itf05g19720; LFC = 4.4975, *p*-value 1.84E-20, 48hr), a putative TF that contains a homeodomain closely linked to a leucine zipper motif, was enriched in both GO terms. In Arabidopsis, ATHB7 and ATHB-12 transcripts were found to be transcriptionally regulated in an ABA-dependent manner and have been shown to mediate drought and respond to ionic osmotic stress (Olsson et al., 2004). Induction of NAC TF genes by drought, salt, and abscisic acid was observed in Arabidopsis (Sakuraba et al., 2015) and the NAC-domain containing TF (itf05g25290: LFC = 2.3674, *p*-value 2.81E-07) was upregulated during salt stress and enriched for response to water and hyperosmotic salinity. Other important genes enriched in salt stress samples were EID1 (itf07g18180: LFC = 2.8208, *p*-value 1.69E-05), an F-box protein involved in phytochrome A-dependent light signal transduction, and fatty acid hydroxylase superfamily (itf04g03470: LFC = 2.8344, *p*-value 1.36E-17), a wax biosynthesis-related gene. It has been demonstrated that water deficit increases the amount of cuticular wax per unit area and leaf cuticle thickness in Arabidopsis plants to enhance their resistance (Lü et al., 2012).

**Figure 4.**
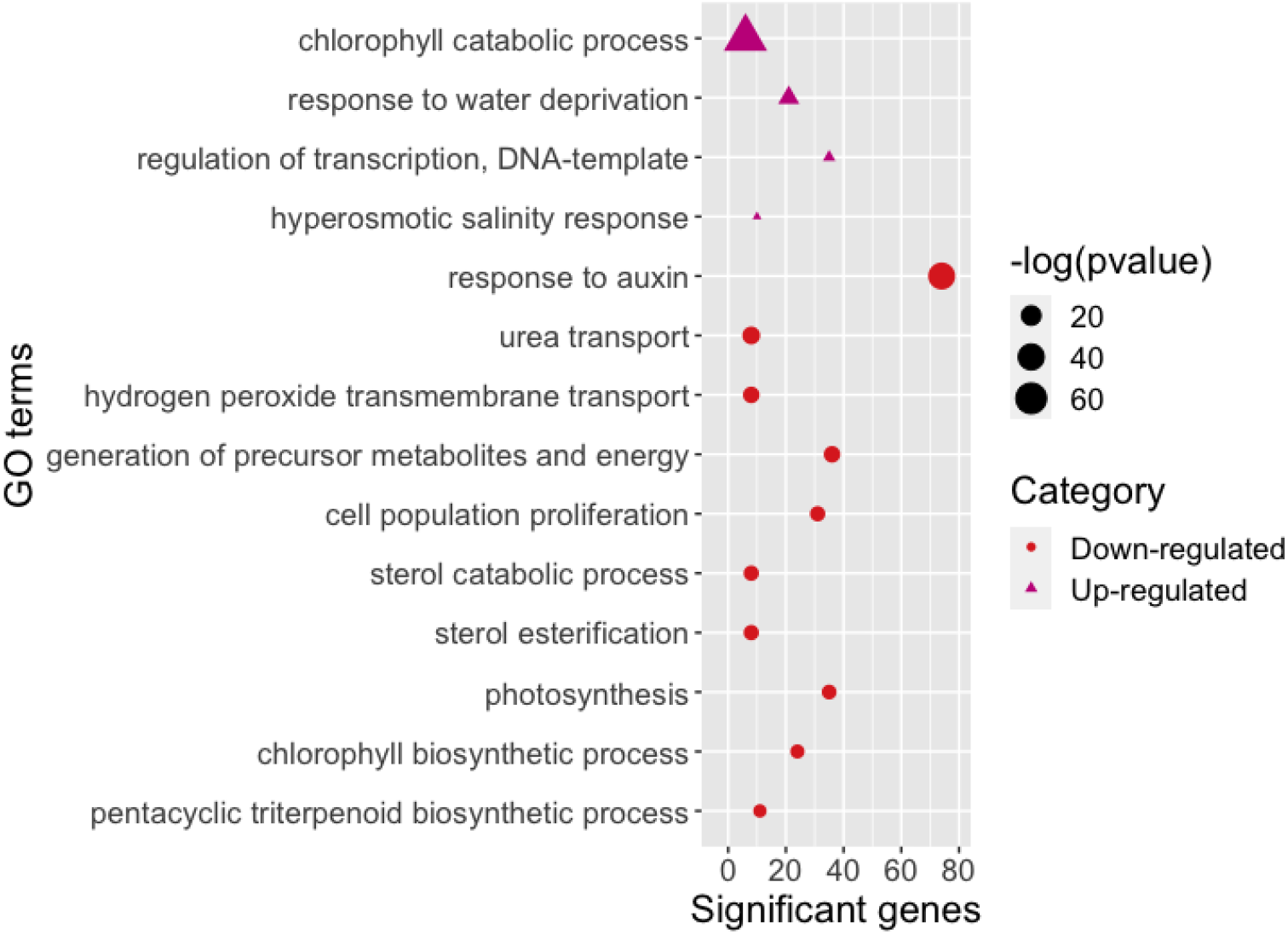
Significant (*p* < 0.05) Gene Ontology (GO) Biological Process terms enriched for upregulated (339; triangles) and downregulated (1108; dots) differentially expressed genes in ‘Beauregard’ leaf tissues at 24 hr and 48 hr of salt stress.

Salt stress resulted in down-regulation of genes associated with ‘chlorophyll biosynthetic process’ (GO:0015995, *p*-value 1.70E-06) and ‘photosynthesis’ (GO:0015979, *p*-value 9.00E-07) (**Table S10)**. Accelerated leaf senescence induced by salt in detached mature sweetpotato leaves treated with NaCl (140mM and 210mM) was accompanied by a reduction in chlorophyll content, reduction of photosynthetic efficiency (Fv/Fm), and an elevation of H_2_O_2_ level (Chen et al., 2012) consistent with the down-regulation of genes involved in photosynthesis observed in this study. Specifically, we observed down-regulation of the light-harvesting in photosystem I and II; NDHK, PHOTOSYSTEM II REACTION CENTER PROTEIN G (PSBG; itf00g02070, LFC = -2.9218, *p*-value 0.003), PHOTOSYSTEM I LIGHT HARVESTING COMPLEX GENE 3 (LHCA3; itf02g14020, 24 hr, LFC = -2.4465, *p*-value 2.21E-10), PHOTOSYSTEM II REACTION CENTER PROTEIN D (PSBD; itf02g22380, LFC = -2.3226, *p*-value, 6.34E-05) and LIGHT HARVESTING COMPLEX PHOTOSYSTEM II (LHCB; itf12g10840, 24 hr, LFC = -2.2464, *p*-value 5.24E-05).

Auxin is involved in the spatial regulation of plant growth and development (Semeradova et al., 2020). Approximately 85 genes encoding auxin and auxin-responsive proteins such as the SAUR-like auxin-responsive protein family were down-regulated following salt stress (**Table S8, S9)**. Multiple SAUR auxin-responsive genes were up-regulated by salt stress including SAUR9 (itf09g00770, 24 hr, log2fc = -3.8313, *p-*value 4.57E-05), SAUR10 (itf12g02790, 24 hr, log2fc = -3.1732, *p-*value 2.57E-07), SAUR3 (itf09g24220, 48 hr, log2fc = -2.8417, *p-*value 0.003; **Table S9**). Hormones including auxin (Hagen and Guilfoyle, 2002; van Mourik et al., 2017) as well as high-temperature (Franklin et al., 2011), drought and high salt conditions (Guo et al., 2018; Wu et al., 2012) can modulate SAUR gene expression and multiple SAUR-like genes were differentially regulated under both heat and salt stress (**Table S9**).

SWEET genes in plants are associated with adaption to adverse environmental conditions including abiotic stress tolerance (Chandran, 2015; Wei et al., 2019; Kafle et al., 2019; Chen et al., 2010) and transcript levels of the Sweet15 gene was reported to be upregulated up to 64-fold higher than the control in Arabidopsis during salt stress (van Zelm et al., 2020; Zhao et al., 2019). Similarly, the expression of MaSWEETs was induced by cold, salt, and osmotic stresses in banana (Miao et al., 2017). In Beauregard, Sugar Will Eventually be Exported Transporters (SWEET) genes (itf08g07850, 24 hr, log2fc = 4.0582, *p-*value 1.05E-09; **Table S5**) was up-regulated following salt stress.

### 3.4 Clustering and co-expression network analyses

Clustering of gene expression patterns is useful in understanding and annotating gene function as ‘guilt-by-association’ can be used to infer gene function. To associate patterns of expression with a stress response, we used the supervised K-means approach and clustered 3,238 of the 3,346 DEGs (after filtering overlapping genes) into 25 clusters (**Figure 5, Table S12**). While most clusters showed no specific pattern associated with stress, clust 15 genes (n = 110) and clust 20 (n = 27) were associated with upregulation during heat stress and enriched for genes associated with heat responses as described above (**Table S13**). Genes that were enriched for clust 15 and 20 included multiple HSPs and HSFs, most of which were upregulated in heat-treated samples but downregulated in salt stress.

**Figure 5.**
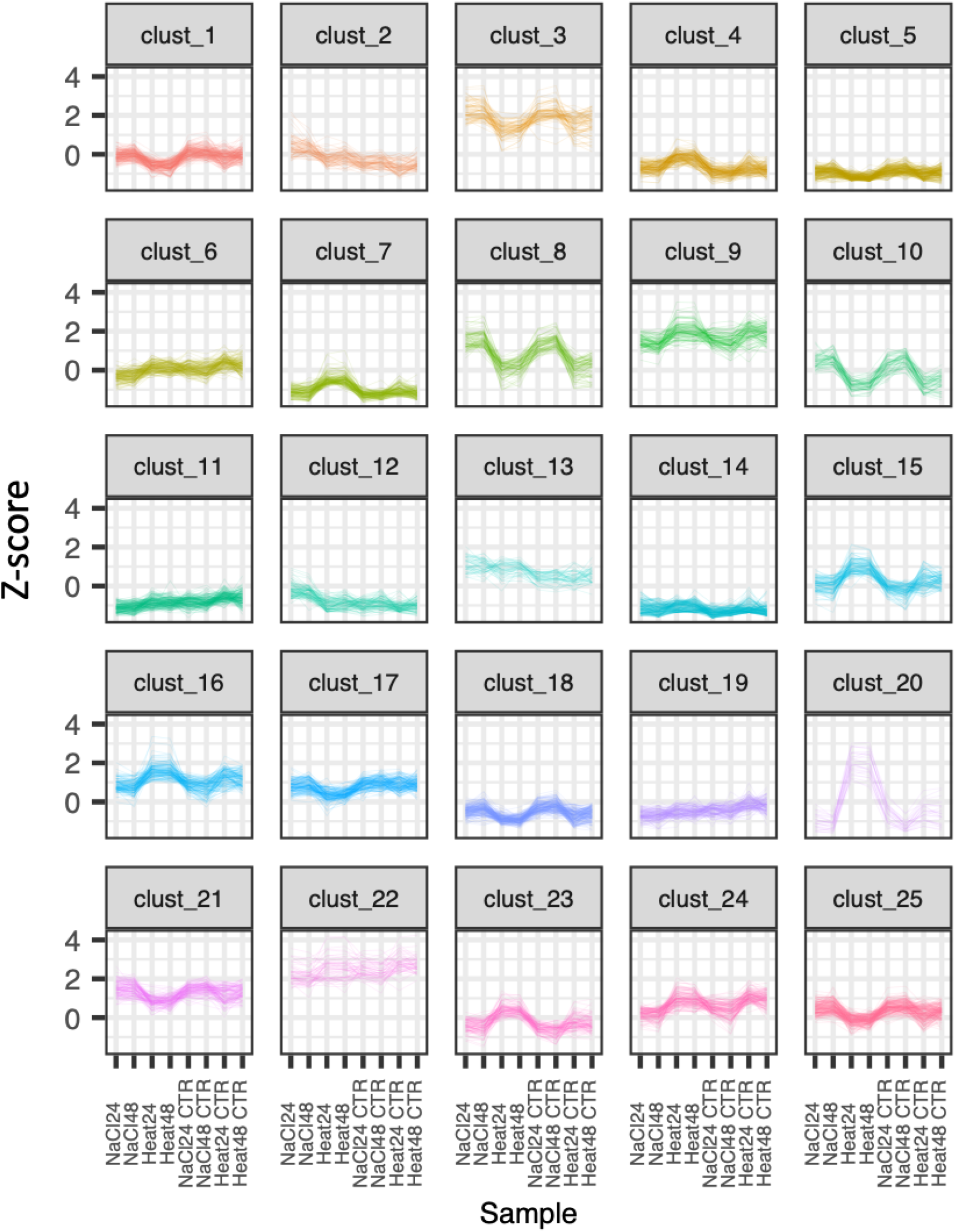
Clustering of gene expression profiles of 3,238 differentially expressed genes by K-means clustering method to identify co-regulated genes following heat and salt stress treatment of Beauregard leaf tissue.

#### 3.4.1 Co-expression network analysis reveals modules of co-regulated genes

Expression abundances (FPKMs) from the heat and salt stress samples were combined with a previous dataset that simulated drought (Lau et al. 2018) to identify genes co-expressed and potentially co-regulated in response to all three stresses. Weighted gene co-expression network analyses identified 18 modules with gene membership ranging from 34 (dark olive green) to 991 genes (salmon) (**Figure S1)** that were enriched in different biological functions (**Table S14**). The blue module was enriched for ‘response to water deprivation’ (GO:0009414, *p*-value; 0.00012), ‘response to abscisic acid’ (GO:0009737, *p*-value; 0.00119), ‘nitrate transport’ (GO:0015706, *p*-value; 2.80E-05), and ‘response to nitrate’ (GO:0010167, *p*-value; 0.00011), among others, suggesting it was associated with drought. Under low N and drought conditions, stomata remain closed; however, combined drought and high N conditions can stimulate stomata to remain open, leading to higher transpiration rates and resulting in greater water use (Shi et al., 2014) thus delaying drought effects (Ren et al., 2015). The HIGH-AFFINITY NITRATE TRANSPORTER 2.7 gene (ATNRT2.7, NRT2; itf03g00630) was up-regulated under both drought (log2fc = 4.6632, *p-value* 2.49E-37; **Table S15**) and salt stress conditions (log2fc = 3.1758, *p-*value 1.20E-11) consistent with previous reports of synergetic cross-talk between N and water transport during water stress (Ding et al., 2018). *AtNRT2*.*7* has been identified as one of the seven high-affinity nitrate transporters in Arabidopsis involved not only in uptake and transport of NO_3_ and its translocation, (Orsel et al., 2006; Wang et al., 2012; Huang et al., 1999) but also in transporting other biological compounds including ABA. Araus et al. (2020) showed that changes in NO_3-_ availability may lead to changes in the expression of drought-responsive genes. Three genes in the blue drought-associated module encoded transcription factors associated with drought stress responses including zf-C2H2 (Cystein2/Histidine; itf11g04060) and HB->HB-HD-ZIP (itf12g07740, itf14g03710) consistent with reports from Arisha et al. (2020b) who found several gene families including C2H2 and HD-ZIP transcription factor family members responding to drought stress in sweetpotato. Transformation of Arabidopsis with *IbZFP, a* C2H2-type zinc finger protein from sweetpotato improved plant drought resistance (Wang et al., 2016) leading to less leaf water loss, lower content of ROS, higher leaf water content, and higher antioxidant enzyme activities after drought treatment of the transgenic plant.

Various TF pathways operating in ABA-dependent and -independent signaling pathways mediate transcriptional regulation of gene expression leading to the expression of early response transcriptional activators and activation of downstream stress tolerance effector genes. The blue drought module contained 63 genes significantly enriched for the GO term ‘response to abscisic acid’ (GO:0009737, *p-*value 0.00119, **Table S14**) with 15 encoding members of multiple TF families. Extensive studies have shown that, in addition to their functions in plant growth and development, NAC transcription factors play a key role in abiotic stress responses (Shao et al., 2015). Transgenic Arabidopsis plants overexpressing either of these paralogs, ANAC019, ANAC055 or ANAC072 (itf05g25290), show significantly increased drought tolerance (Tran et al., 2004). In our samples, the NAC gene (itf05g25290) was present in the drought-associated module (blue) and was differentially upregulated during drought and salt stress (itf05g25290, salt; 48 hr, LFC = 2.3674, *p*-value 2.81E-07, **Table S5**; drought; 48 hr, LFC = 7.0490, *p*-value 1.83E-53; **Table S15**). In Arabidopsis, ARABIDOPSIS THALIANA HOMEOBOX 7 (ATHB-7) and AtHB12 genes are strongly up-regulated after osmotic or drought stresses in young plants upon ABA or NaCl treatment (Lee and Chun, 1998; Olsson et al., 2004; Söderman et al., 1996). Furthermore, ectopic expression of AtHB7 confers drought tolerance to transgenic tomato (Mishra et al., 2012). The homeobox gene itf05g19720 was upregulated in response to salt (LFC = 3.7014, *p-*value 1.84E-20) and drought (LFC = 8.9468, *p*-value 1.20E-86) at 48 hr timepoint in our study.

#### 3.4.2 Key hub genes associated with drought sensitivity in sweetpotato

Within the drought-associated (blue) module, osmotin (itf13g03220, LFC = 5.9807, *p*-value 8.67E-108; 48 hr) was the highest connected (*ktotal* and *kwithin;* **Figure 6; Table S16)**. Osmotin, a multifunctional stress-responsive protein, belongs to the pathogenesis-related 5 (PR-5) defense-related protein family and imparts drought, salt, and cold tolerance (Goel et al., 2010; Husaini and Abdin, 2008; Le et al., 2018; Patade et al., 2013; Weber et al., 2014; Wan et al., 2017; Zhu et al., 1995). During stress conditions, the accumulation of the osmolyte proline which quenches ROS and free radicals is facilitated by osmotin. Furthermore, osmotin from the resurrection plant *Tripogon loliiformis* confers numerous simultaneous abiotic stresses tolerance (cold, drought, and salinity) in transgenic rice (Le et al., 2018; Mandal et al., 2018). Overexpression of the osmotin gene in early stages of osmotic stresses (cold, drought, and salinity) by a 1000-fold conferred resistance against abiotic stress in transgenic *Nicotiana tabacum, Oryza sativa*, and sesame (Chowdhury et al., 2017) while in potato, osmotin over expression was found to cause delays in the development of late blight disease symptoms (Liu et al., 1994). As a result, osmotin has been proposed as a high-value gene for developing multiple stress-tolerant biofortified crops (Husaini, 2022).

**Figure 6.**
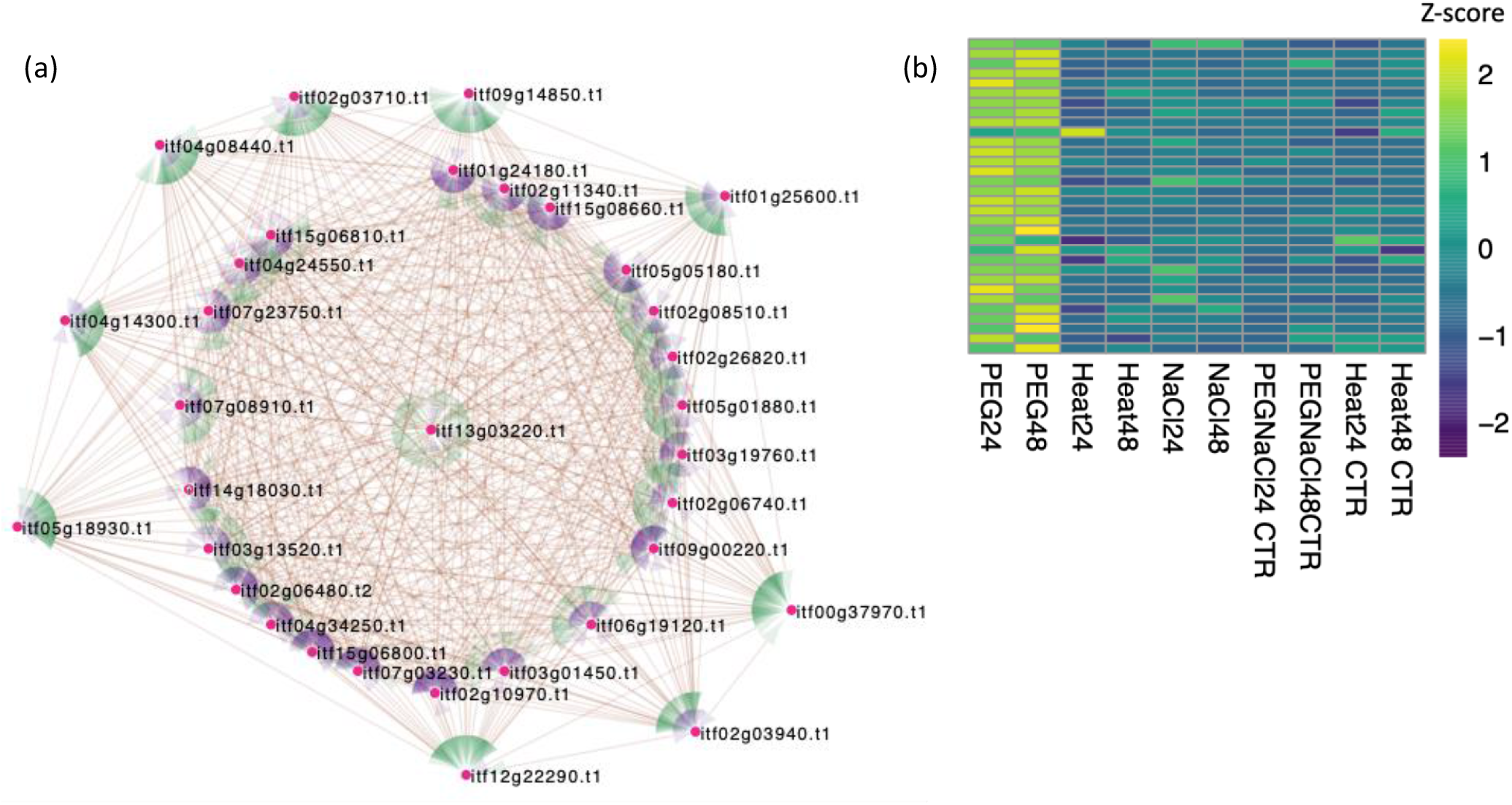
Weighted gene co-expression network analysis. (WGCNA). (a) Coexpression network including 32 highly connected nodes (genes, magenta) from the drought (blue) module. Targeted and source nodes are indicated by green and purple colors, respectively. (b) Heatmap displaying expression (Z-score) of (blue) drought network genes from heat, salt (NaCl) and drought (PEG) treated Beauregard, leaf tissue.

Among the terms enriched for the blue, drought module, were genes in response to ‘extracellular region’ (GO:0005576, *p*-value; 4.50E-14). Furthermore, two expansin A15 genes found in our network analysis were up-regulated in the drought-stressed samples; itf02g03710 (LFC = 9.3730, *p*-value 5.14E-97) and itf02g03940 (LFC = 9.7680, *p*-value 3.87E-79) at 48hr.

Expansins are induced by phytohormones, biotic and abiotic stresses such as heat, drought, salt and heavy metals and are involved in cell expansion and cell wall changes (Cosgrove, 2000; Ding et al., 2016; Lu et al., 2016; Le Gall et al., 2015; Xu et al., 2014). It has been shown that the overexpression of expansin genes in *Arabidopsis* enhances plant tolerance to drought stress, salt stress (Dai et al., 2012), tolerance to drought in transgenic tobacco, (Li et al., 2011), high salt in wheat (Han et al., 2012) and rice (Jadamba et al., 2020), and drought and heat tolerance in potato (Chen et al., 2019). Interestingly, besides drought and salt having common response patterns, osmotin and expansin A15 genes were not differentially expressed in salt stress samples which may suggest the presence of specifically affected biochemical pathways by drought.

## 4.0 Conclusions

Characterization of heat, salt, and drought-responsive gene expression in an orange-fleshed sweetpotato cultivar identified stress specific as well as shared responses to the stress treatments. A key impact of abiotic stress was evident from the downregulation of photosynthesis, arrest of the cell cycle, and activation of a canonical heat shock response. We identified crosstalk signaling between genes, TFs and phytohormones. Construction of co-expression modules delineated TFs and hub genes of interest for future work including improving abiotic stress tolerance in sweetpotato.

## Author’s contributions

DCG and AK designed and executed the experiments and isolated RNA. SW and ZF sequenced the RNA for gene expression profiling. MK performed RNAseq bioinformatics analysis guided by JW, JH and CRB. Data visualization and manuscript writing was performed by MK and CRB. All authors read and approved the manuscript.

## Supplementary Materials

### Supplementary Figures

**Figure S1.**
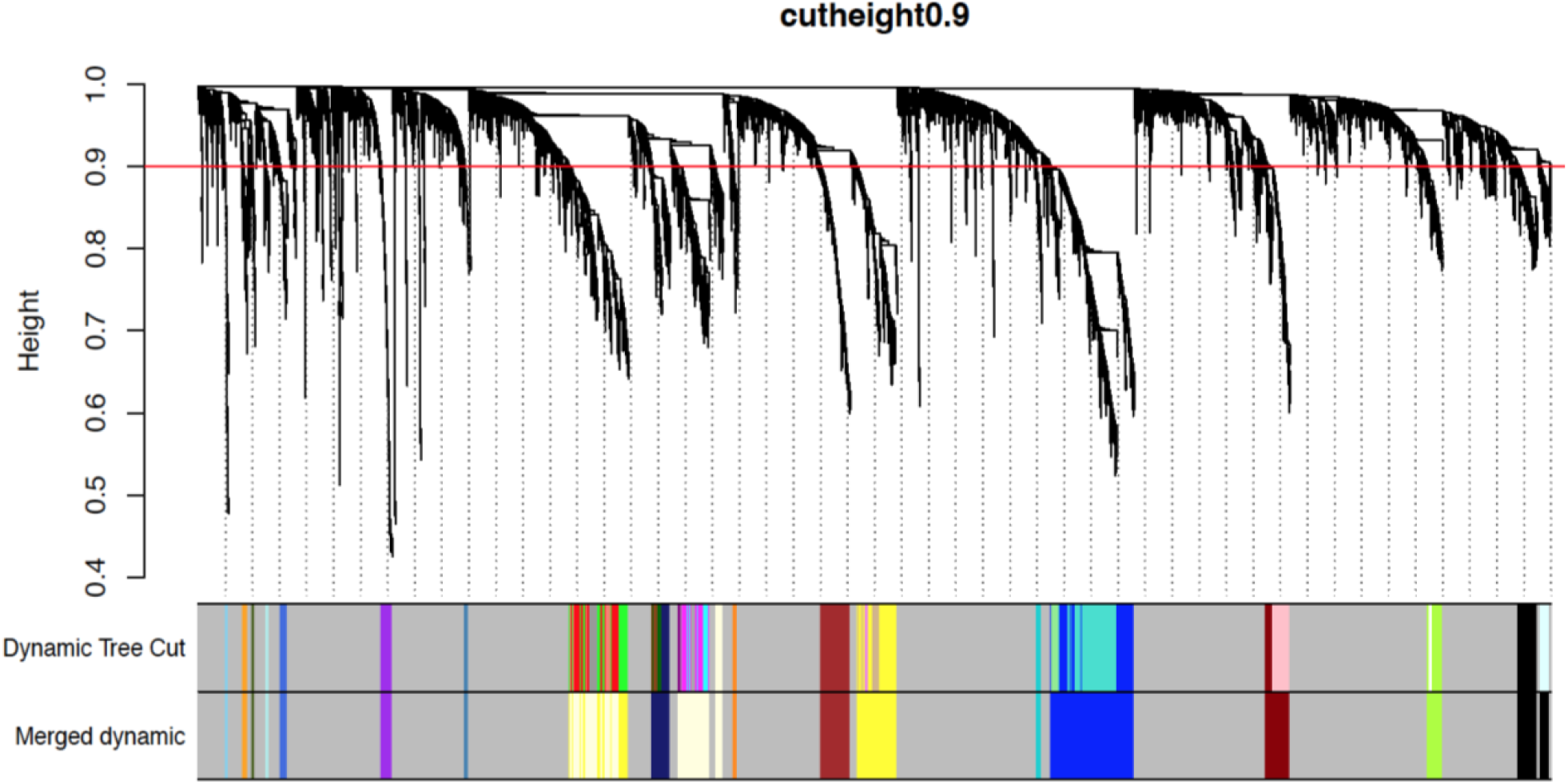
WGCNA clustering dendrogram of genes obtained by hierarchical clustering of adjacency-based dissimilarity of 14,138 genes from the heat, salt and drought treated Beauregard leaf tissue. Coexpression modules were identified *via* the Dynamic Tree Cut method; the merged dynamic indicates modules divided according to similarity of the module (with assigned module colors). Analysis was carried out according to the merged modules. Vertical distance in tree diagram represents distance between two nodes (between genes)

### Supplementary Tables

**Table S1**. Summary of RNA-Seq libraries and alignment metrics.

**Table S2**. Pearson’s Correlation Coefficients of control, heat- and salt-stress RNA-seq libraries.

**Table S3**. Differential expression of all up-regulated (LFC > 2.0) and downregulated (LFC < 2.0) genes at 24 hr and 48 hr stress following heat stress and salt stress treatments of Beauregard.

**Table S4**. Unique differentially up-regulated genes following heat stress (DEGs; LFC > 2.0; p-value < 0.05).

**Table S5**. Unique differentially up-regulated genes following salt stress (DEGs; LFC > 2.0; p-value < 0.05).

**Table S6**. Common differentially up-regulated genes following heat and salt stress (DEGs; LFC > - 2.0; p-value < 0.05).

**Table S7**. Unique differentially down-regulated genes following heat stress (DEGs; LFC < - 2.0; p-value < 0.05).

**Table S8**. Unique differentially down-regulated genes following salt stress (DEGs; LFC < - 2.0; p-value < 0.05).

**Table S9**. Common differentially down-regulated genes following heat and salt stress (DEGs; LFC > - 2.0; p-value < 0.05).

**Table S10**. Gene ontology (GO) terms enriched for the up- and down differentially regulated genes following heat and salt stress treatments on Beauregard sweetpotato leaf tissues at 24 and 48 hours.

**Table S11**. Expression (gene counts as log CPM) of heat shock-related genes in heat and salt treated Beauregard sweetpotato leaf samples.

**Table S12**. Differentially regulated genes grouped into 25 clusters using K-means based on the expression patterns (Z-score) in the treated and control samples with transcription factor information.

**Table S13**. Gene ontology (GO) term enrichment of genes in K-means clusters.

**Table S14**. Gene ontology enrichment of Weighted Gene Coexpression Network Analysis (WGCNA) modules.

**Table S15**. Differentially up-(LFC > 2.0) and downregulated (LFC < 2.0) genes following drought (25% PEG) stress; p-value < 0.05).

**Table S16**. Intra (*kwithin*) and inter (*ktotal*) modular connectivity measuring of genes in the Weighted Gene Coexpression Network Analysis.

